# LINE-2 transposable elements are a source for functional human microRNAs and target sites

**DOI:** 10.1101/218842

**Authors:** Rebecca Petri, Per Ludvik Brattås, Yogita Sharma, Marie E Jönsson, Karolina Pircs, Johan Bengzon, Johan Jakobsson

## Abstract

Transposable elements (TEs) are dynamically expressed at high levels in multiple human tissues, but the function of TE-derived transcripts remains largely unknown. In this study, we identify numerous TE-derived microRNAs (miRNAs) by conducting Argonaute2 RNA Immunoprecipitation followed by small RNA sequencing (AGO2 RIP-seq) on human brain tissue. Many of these miRNAs originated from LINE-2 (L2) elements, which entered the human genome around 100-300 million years ago. We found that L2-miRNAs derive from the 3’ end of the L2 consensus sequence and thus share very similar sequences, indicating that they could target transcripts with L2s in their 3’UTR. In line with this, we found that many protein-coding genes carry fragments of L2-derived sequences in their 3’UTR, which serve as target sites for L2-miRNAs. L2-miRNAs and targets were generally ubiquitously expressed at low levels in multiple human tissues, suggesting a role for this network in buffering transcriptional levels of housekeeping genes. Interestingly, we also found evidence that this network is perturbed in glioblastoma. In summary, our findings uncover a TE-based post-transcriptional network that shapes transcriptional regulation in human cells.

## Introduction

The emergence and evolution of gene regulatory networks is thought to underlie biological adaptations and speciation. Transposable elements (TEs) have been implicated in these processes since they can amplify in numbers and move into new regions of the genome. Genomic analyses supports a role for TEs in gene regulatory networks, since a substantial fraction of TEs evolve under selective constraints despite being non-coding (Chuong et al. 2017) but the actual impact of TEs on human transcriptional networks remains poorly understood.

We and others have recently described that many transcripts expressed in various human tissues contain TE-derived sequences (Brattas et al. 2017; Deininger et al. 2017). Many of these sequences appear to be indirectly transcribed, often in antisense direction, as part of other transcripts including those coding for protein (Deininger et al. 2017). Together with the finding that TEs can generate miRNA target sites in the 3’UTR of protein-coding transcripts (Shin et al. 2010; Spengler et al. 2014) the data suggests that TEs within transcripts can at least in part act as templates for RNA-binding proteins such as the microRNA (miRNA) machinery and contribute to post-transcriptional regulation.

In addition, TEs can be a source for non-coding RNAs, such as long non-coding RNAs (lncRNAs) or miRNAs. Computational and experimental studies have shown that LINE (e.g. LINE-1, LINE-2, LINE-3), SINE (e.g. MIR and Alu) and some LTR containing transposons can act as miRNA sources (Piriyapongsa et al. 2007; Roberts et al. 2014; Spengler et al. 2014). Functionally validated TE-derived miRNAs are for instance miR-28, miR-95 and miR-151 that are derived from LINE-2 (L2) elements (Ding et al. 2010; Shin et al. 2010; Frankel et al. 2014). However, current studies demonstrating functional TE-derived miRNAs and target sites are mainly conducted in cell lines and functional evidence for a TE-based post-transcriptional network in primary tissues, such as the human brain, is missing.

In this study, we used Argonaute2 RNA Immunoprecipitation (AGO2 RIP) on adult human brain tissue. We found that many small RNAs bound by AGO2 are derived from TEs, with the majority being derived from L2 elements. These L2-miRNAs show strong sequence complementarity to L2 elements found in the 3’UTR of protein coding genes. Transcripts containing L2 elements in the 3’UTR are incorporated into the RNA induced Silencing complex (RISC) in the human brain and are regulated by L2-miRNAs. We found that L2-miRNAs and targets are generally ubiquitously expressed in multiple human tissues, hereby primarily acting on housekeeping genes. We also found that this L2-miRNA network is perturbed in samples from patients with glioblastoma. Together our results demonstrate a TE-based post-transcriptional network that influences the expression of protein-coding genes in human tissues.

## Results

### Identification of transposable element-derived microRNAs expressed in the human brain

To identify small RNAs that participate in gene silencing in the human brain we performed Argonaute2 RNA interacting Immunoprecipitation followed by small RNA sequencing (AGO2 RIP-seq) on surgical biopsies obtained from human cortex (n = 3) (Fig 1A). The use of AGO2 RIP-seq on fresh human brain tissue circumvents several challenges associated with detecting functional small RNAs derived from transposable elements (TEs) since it reduces background noise generated by degradation products as well as avoids problems arising with the use of cell lines, where a loss of DNA methylation could activate aberrant TE expression.

**Figure 1:**
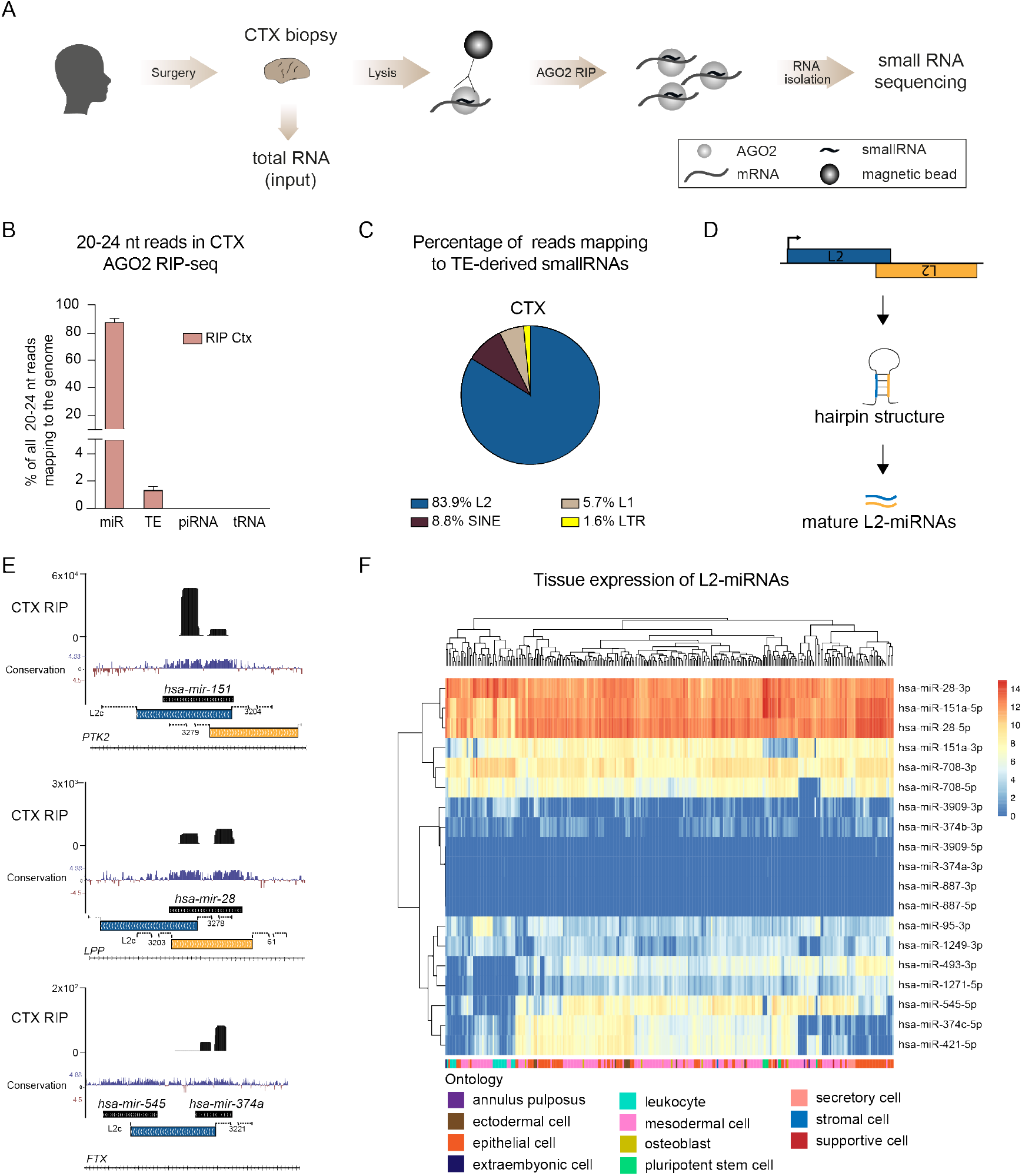
AGO2-associated small RNAs in human cortex tissue. A) Schematics of AGO2 RIP-seq on human cortex samples followed by small RNA sequencing. B) Bar graph showing the percentage of 20-24 nucleotide (nt) long reads in the human genome mapping to mature miRNAs (miR), transposable elements (TE), piwi RNAs (piRNA) and transfer RNAs (tRNA). Data is represented as mean ± SEM (RIP Ctx n = 3). C) Pie chart showing the percentage of reads mapping to TEs. D) Schematics of the generation of miRNAs from L2 elements. E) UCSC genome browser tracks showing examples of L2-miRNAs in AGO2 RIP-seq samples from human brain tissue. F) Heatmap depicting the expression of L2-miRNAs across different human tissues and cell types (n = 400). AGO – Argonaute, AGO RIP – AGO RNA Immunoprecipitation, CTX – cortex, L1 – LINE1, L2 – LINE-2, LTR – long terminal repeat, mir – precursor miRNA, SINE – short interspersed nuclear element.

We found that AGO2-bound small RNAs displayed a high enrichment for 20-24 nucleotide (nt) long reads, the typical size of microRNAs (miRNAs), while input samples, which include all small RNAs in the tissue, displayed an expected broad size profile of RNAs including e.g. many RNAs in the size range of 30-36 nt (S1A Fig). We next investigated the genomic origin of small RNAs expressed in the human brain. As expected, most AGO2-bound RNAs were classical miRNAs (S1B Fig) and we found very limited evidence for AGO2-binding of transfer RNA fragments (tRFs) or piwi-RNAs (piRNA), although tRFs, of mostly 30-36 nt in size, were abundant in input samples (S1B Fig).

When focusing our analyses on AGO2-bound small RNAs of 20-24 nt in size we detected high expression of classic brain-enriched miRNAs, such as miR-128 (S1C Fig) (Tan et al. 2013). Interestingly, we also found a substantial fraction (around 2-3 % of reads) of AGO2-bound RNAs that mapped to TEs, including LINE, LTR and SINE elements (Fig 1B).

### L2-microRNAs are abundantly expressed in human tissues

To identify miRNA-like small RNAs that are derived from TEs, we aligned reads with a length of 20-24 nt to the human reference genome (hg38) and removed all reads that mapped equally well to more than one locus (see methods for a more detailed description). Surprisingly, the great majority (83.9 %) of the identified TE-derived small RNAs originated from L2 elements (Fig 1C). When comparing the L2-derived small RNAs with miRbase annotations, we found that they have previously been identified as miRNAs including e.g. L2c-derived miR-151 and L2b-derived miR-95 (Table 1) (Smalheiser and Torvik 2005; Piriyapongsa et al. 2007). Many of the L2-miRNAs originate from two distinct L2 elements that are located in close vicinity and oriented in opposite direction to each other, thereby providing a source of hairpin structures (Fig 1D, E) (Smalheiser and Torvik 2005). However, we also found cases where a single L2 element gives rise to two miRNA precursors e.g. *mir-545* and *mir-374* (Fig 1E).

**Table 1:** Expressed L2-derived miRNAs in human brain tissue.

| TE | coordinates | direction of<br>microRNA | miRNA | paired | mean expression<br>(reads in RIP) |
| --- | --- | --- | --- | --- | --- |
| L2b | chr 4: 8005199 - 8005343 (+) | AS | miR-95-3p | L2c | 8344.66 |
| L2c | chr 8: 140732529 - 140732650 (-) | S | miR-151a-3p/5p | L2c | 5253.32 |
| L2c | chr 8: 140732622 - 140732734 (+) | AS | miR-151a-5p | L2c | 29.13 |
| L2c | chr 3: 188688700 - 188688813 (-) | AS | miR-28-5p | L2c | 771.25 |
| L2c | chr 3: 188688783 - 188688877 (+) | S | miR-28-3p/-5p | L2c | 299.29 |
| L2c | chr 11: 79402022 - 79402117 (+) | AS | miR-708-3p/-5p | L2c | 695.05 |
| L2c | chr 11: 79401986 - 79402051 (-) | S | miR-708-3p | L2c | 217.33 |
| L2c | chr 22: 45200672 - 45200995 (-) | S | miR-1249-3p | - | 443.35 |
| L2c | chr X: 74287157 - 74287324 (-) | S | miR-374a-3p/ miR-545-5p | - | 90.09 |
| L2c | chr X: 74218419 - 74218583 (-) | S | miR-374b-3p/ miR-421 5p | - | 12.58 |
| L2c | chr X: 74218419 - 74218583 (-) | AS | miR-374c-5p | - | 12.58 |
| L2b | chr14: 100869090 - 100869150 (-) | S | miR-493-3p |  | 49.70 |
| L2b | chr 5: 176367379 - 176367999 (+) | S | miR-1271-5p | L2a | 31.11 |
| L2c | chr 22: 35335672 - 35335822 (-) | AS | miR-3909-3p/ -5p | - | 71.28 |
| L2a | chr 5: 15934793 - 15935362 (-) | AS | miR-887-3p/5p | - | 9.34 |

To investigate the similarity between different L2-miRNAs we focused on those deriving from the L2c-subfamily and mapped all detected L2c-miRNAs to the L2c consensus sequence (RepeatMasker). This analysis showed that many L2c-miRNAs are similar in sequence since they are generated from the same position within the 3’end of the L2c consensus with only a few bases difference, therefore most likely sharing many targets (S1D Fig).

In total, we detected 19 high-confidence L2-miRNAs expressed in the human brain (Table 1, Fig 1F). To investigate the expression of these L2-miRNAs in other human tissues we used a publicly available smallRNA-seq dataset from 400 human samples (de Rie et al. 2017). We found that most L2-miRNAs are ubiquitously expressed, albeit at different levels, including high-level expression of miR-28-3p, miR-28-5p and miR-151a-5p (Fig 1F). On the contrary, we found very limited evidence for cell-type specific expression of L2-miRNAs (Fig 1F).

### L2-derived miRNAs are expressed in the mouse brain

To investigate whether L2-miRNAs are conserved among mammals, we also conducted AGO2 RIP-seq on mouse brain tissue (Fig 2A). We found a similar proportion of TE-derived miRNAs in the mouse brain as in the human brain, with L2-miRNAs being the most common TE-derived miRNAs also in mouse (Fig 2B-C, Table 2). Several of the L2-miRNAs were conserved between mouse and human including e.g. miR-151 (Fig 2D, Table 1&2). However, we also noted that some L2-miRNAs were present in the human but not mouse genome, e.g. miR-95 (Table 1&2). Additionally, we identified miRNAs such as hsa-miR-28 and mmu-miR-28c, with the same mature sequence in the human and mouse genome, however, originating from different unique retrotransposition events (Fig 2D, Table 1&2). This shows that L2-miRNAs are bound by AGO2 and expressed in the brain of different mammalian species, suggesting a conserved functional role for these non-coding RNAs. However, it is also worth noting that we detected a higher proportion of miRNAs derived from other TEs such as LINE-1 (L1) and MIRs in the mouse brain compared to human samples.

**Figure 2:**
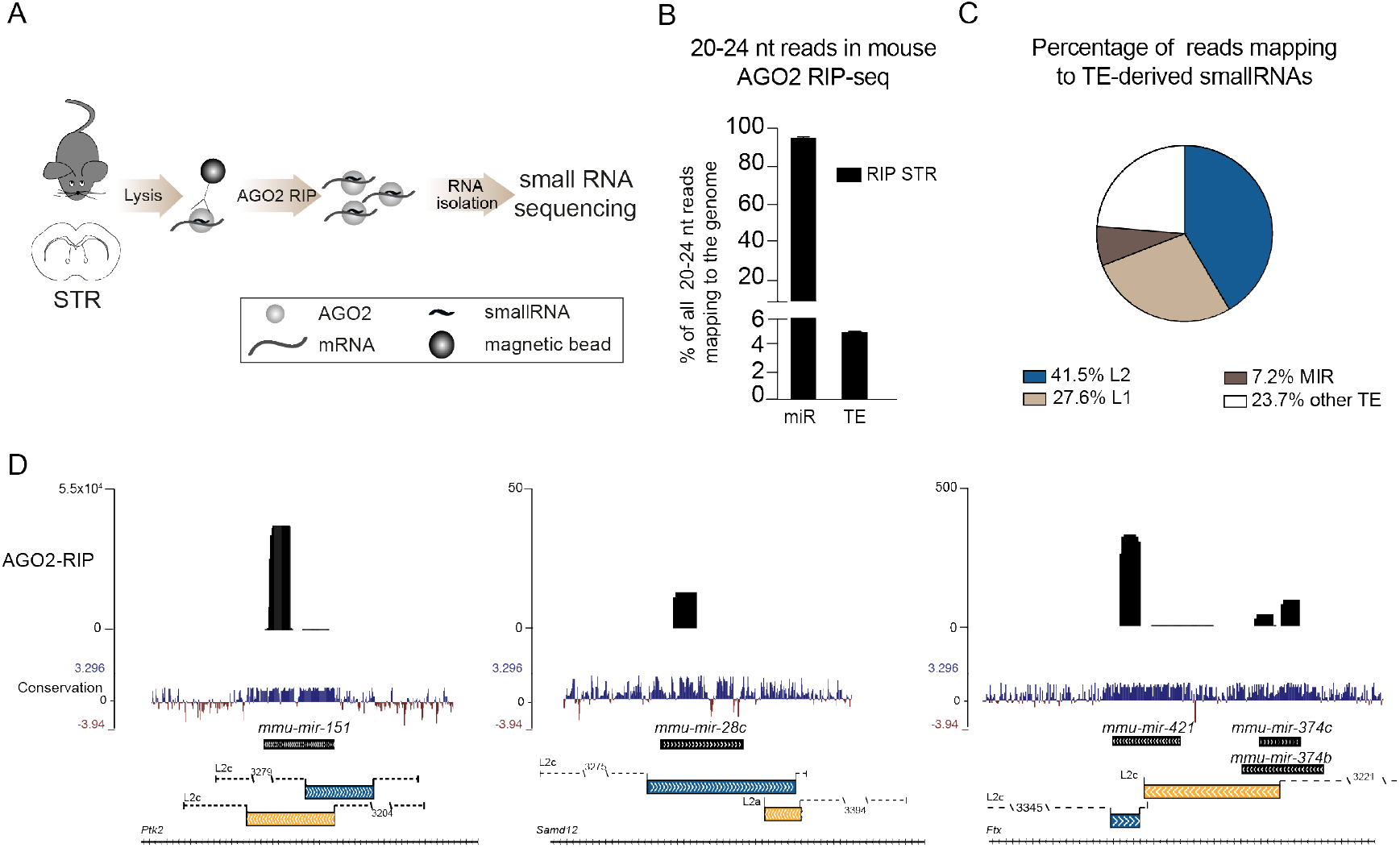
AGO2-associated small RNAs in the mouse brain. A) Schematics of AGO2 RIP-seq on mouse striatum (STR) followed by small RNA sequencing. B) Bar graph showing the percentage of 20-24 nt long reads in the mouse genome mapping to mature miRNAs (miR) and transposable elements (TE). Data is represented as mean ± SEM (RIP STR n = 3). C) Pie chart showing the percentage of reads mapping to TEs. D) UCSC genome browser tracks showing examples of L2-miRNAs in AGO2 RIP-seq samples from mouse brain tissue. AGO2 – Argonaute2, AGO RIP – AGO-RNA Immunoprecipitation, L1-LINE-1, L2 – LINE-2, mir – precursor miRNA, MIR - Mammalian-wide interspersed repeat.

**Table 2:** L2-derived miRNAs expressed in mouse brain tissue.

| TE | coordinates | direction of<br>miRNA | miRNA | paired | mean expression<br>(reads in RIP) |
| --- | --- | --- | --- | --- | --- |
| L2c | chr 15: 73254797 - 73254856 (-) | S | miR-151-3p/5p | L2c | 41322 |
| L2c | chr 15: 73254854 - 73254920 (+) | AS | miR-151-5p | L2c | 6 |
| L2c | chr 7: 96249497 - 96249546 (+) | S | miR-708-3p | L2a | 25417 |
| L2a | chr 7: 96249441 - 96249478 (-) | AS | miR-708-5p | L2c | 183 |
| L2c | chr 4: 94665178 - 94665336 (-) | S | miR-872-3p/5p | - | 2965 |
| L2c | chr 15: 84951302 - 84951582 (-) | S | miR-1249-3p | - | 2064 |
| L2c | chr 16: 24827889 - 24827951 (+) | S | miR-28a-3p | L2c | 392 |
| L2c | chr 16: 24827826 - 24827887 (-) | AS | miR-28a-5p | L2c | 65 |
| L2c | chr X: 103572919 - 103572951 (+) | AS | miR-421-3p | L2c | 284 |
| L2c | chr X: 103572957 - 103573111 (-) | S | miR-421-5p/ miR-374b-3p | L2c | 37 |
| L2c | chr X: 103572957 - 103573111 (-) | AS | miR-374c-5p | L2c | 37 |
| L2b | chr 12: 109580258 - 109580318 (-) | AS | miR-493-3p/5p | - | 161 |
| L2a | chr X: 105379137 - 105379236 (-) | S | miR-325-5p | L2b | 110 |
| L2b | chr X: 105378963 - 105379128 (+) | AS | miR-325-3p | L2a | 40 |
| L2 | chr 10: 128382803 - 128382885 (-) | AS | miR-6914-3p/5p | - | 41 |
| L2a | chr X: 147010598 - 147010647 (+) | S | miR-1264-5p | - | 29 |
| L2c | chr15:53614194 - 53614328 (+) | AS | mir-28c-3p/5p | - | 4 |

### L2-derived fragments are found in the 3’UTR of protein coding genes

L2-derived AGO2-associated miRNAs have a large number of potential “self-targets” since these elements extensively colonized the genome of our ancestors around 100-300 million years ago, resulting in almost 500,000 L2-derived fragments in the human genome (Kapitonov 2006). Thus, L2-miRNAs could potentially guide the RISC to 3’UTRs containing these elements transcribed in the opposite direction of their element of origin, thereby providing a possibility for TEs to post-transcriptionally influence the expression of numerous protein-coding genes. To investigate this possibility, we analysed the location of L2 elements in the human genome and found that a substantial number of L2 elements, 2847, are located in 3’UTRs thereby having the potential to act as miRNA-targets.

To investigate the expression profile of these genes in various human tissues we used a publicly available data set containing RNA-seq from 27 different human tissues and cells (Fagerberg et al. 2014). We found that the majority of genes that carry L2s in the 3’UTR are ubiquitously expressed, mostly at low levels (Fig 3A). Using the classification for tissue specific expression from Fagerberg et al. (Fagerberg et al. 2014), we found that the vast majority of these genes are expressed in all tissues at low levels (Fig 3B). On the contrary, only a small proportion of these genes display tissue-specific expression (Fig 3B).

**Figure 3:**
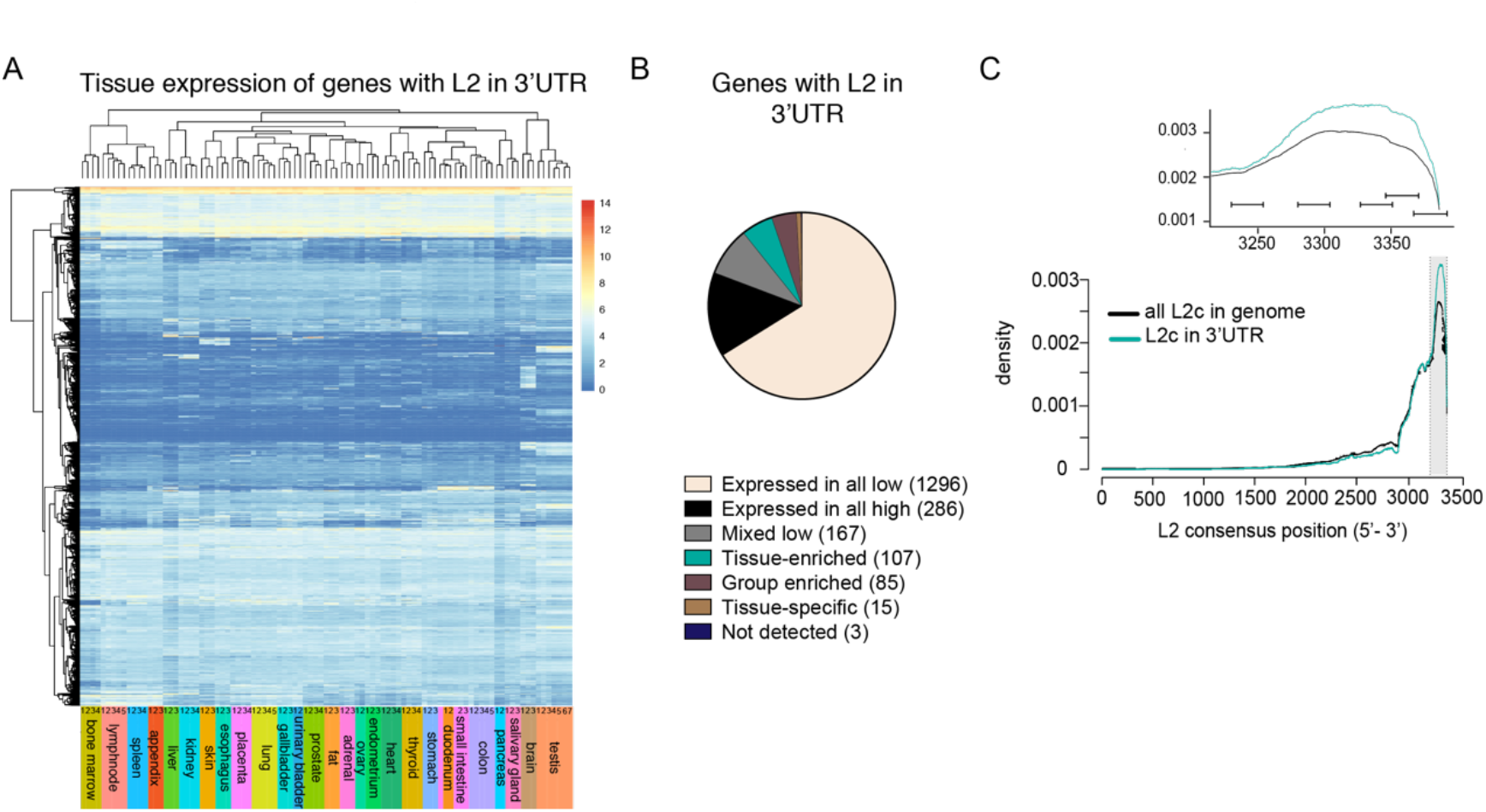
Expression of L2-carrying genes in human tissues. A) Heatmap depicting the expression of L2-carrying genes across 27 tissue samples obtained from 95 individuals (Fagerberg et al. 2014). B) Pie chart showing the number of expressed genes classified based on the Fagerberg tissue specificity classification (Fagerberg et al. 2014). C) Graph showing the density of the L2c consensus sequence of all L2c in the genome (black line) and of L2c element in the 3 ‘UTR (blue line).

We next analysed the structural conservation of L2 elements in the human genome, focusing on the L2c family. We found that the majority of L2c are 5’-truncated, as it is often the case for LINE-elements. Interestingly, when analysing specifically L2c elements located in the 3’UTR of protein coding genes, we found an even stronger conservation of the very 3’-end of the L2 element, which corresponds to the region from where the L2-miRNAs are generated (Fig 3C). Together, these data demonstrate that around 2000 genes, mostly lowly expressed housekeeping genes, carry a L2 element in the 3’UTR. These L2 elements have the potential to act as target sites for the RISC, since they are conserved to match the sequence of the L2-miRNAs.

### L2-derived fragments in the 3’UTR of protein coding genes are L2-microRNA-targets

To provide functional evidence that L2-containing 3’UTRs are regulated by miRNAs in the human brain, we conducted AGO2 RIP on cortex tissue (n = 2) followed by total RNA sequencing to identify miRNA target genes (Fig 4A). As expected, most reads in the RIP-samples were mapping to RefSeq annotations (Fig 4B), while input samples were rich on ribosomal RNA (rRNA). The amount of rRNA was, as expected, strongly decreased in the RIP-samples (Fig 4B). We found an enrichment of reads mapping to 3’UTRs of genes in the RIP samples compared to input fractions, which is line with the high prevalence of miRNA target sites in this part of a transcript (Fig 4B). Strikingly we also found that reads mapping to TEs were enriched in RIP-samples.

**Figure 4:**
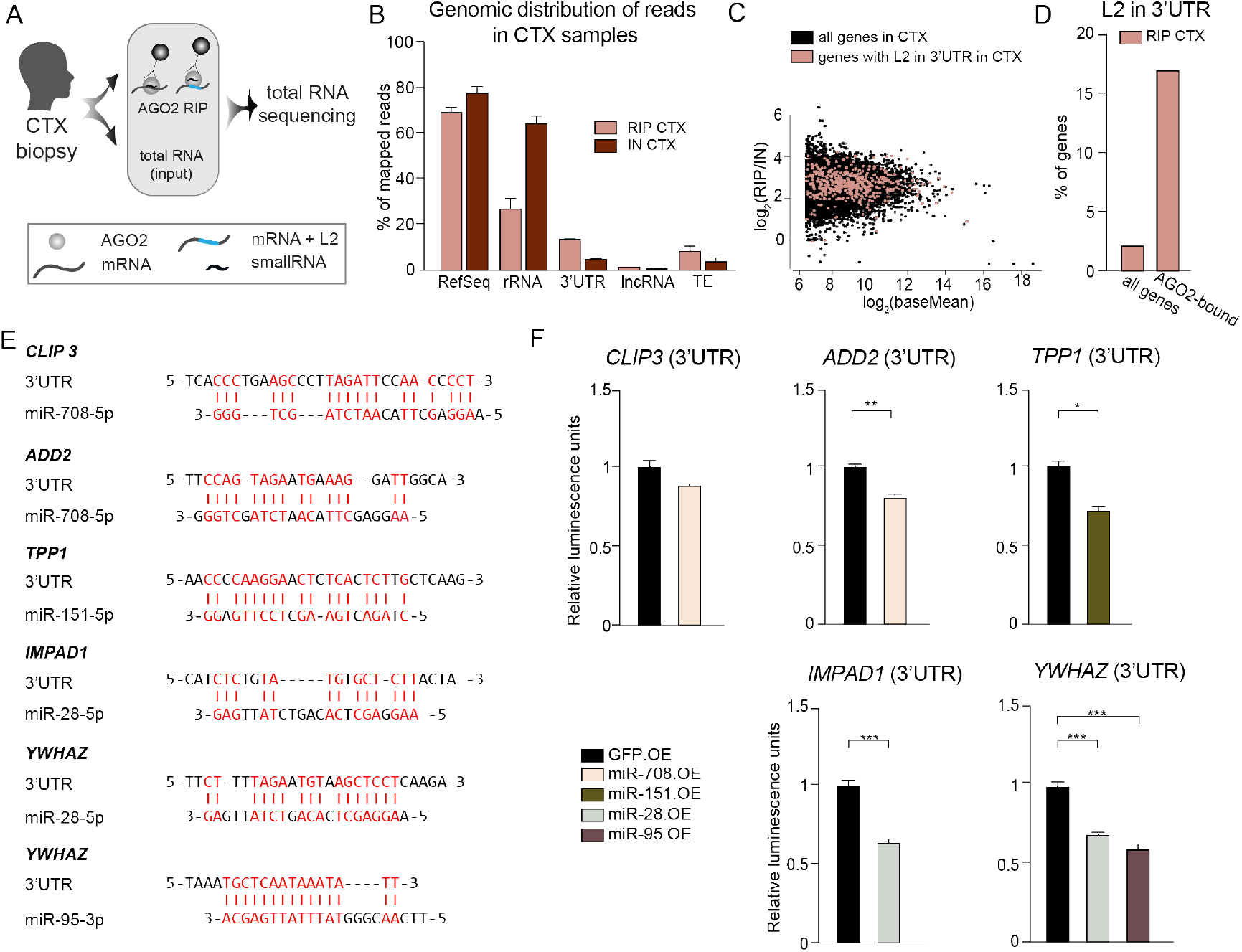
Genes with L2 in 3’UTR are bound by AGO2 in the human brain. A) Schematics of AGO2 RIP followed by total RNA sequencing on cortex (CTX) biopsies. B) Genomic distribution of RNAs in the AGO2 RIP-seq and input samples of cortex (CTX, n = 2). C) Dot plot of all genes in cortex samples of the human brain. Genes with L2 in the 3 ‘UTR are marked in red. The log2 transformed mean expression (x-axis) is plotted against the log2 transformed fold change of RIP (n = 2) versus input samples (n = 2) (y-axis). D) Bar plot showing the percentage of genes with in the 3’UTR of all genes, and of the top 100 highest expressed genes that are more than 4-fold enriched in RIP compared to input samples. E) Potential target sites of L2-miRNAs in the L2 sequence in 3 ‘UTRs of genes. Complementary bases are marked in red and with a vertical line. F) Luciferase assay of L2-derived target sites. Data is shown as mean ± SEM. AGO-Argonaute, AGO2 RIP – AGO2 RNA Immunoprecipitation, CTX – Cortex, L2 – LINE-2, lncRNA – long non-coding RNA, OE – overexpressor, TE – transposable elements, rRNA – ribosomal RNA.

We next analysed all genes with L2 elements in their 3’UTR (Fig 4C) and set stringent criteria to identify high-confidence AGO2-bound genes. We selected the 100 most abundant transcripts in the RIP samples, which were also more than 4-fold enriched in RIP compared to the input fractions. Strikingly, we found that among the AGO2-bound genes, L2-carrying transcripts were highly enriched and made up 15 % of the transcripts (Fig 4D). In comparison, approximately 2 % of all transcripts detected in this analysis carry L2 in their 3’UTR. This analysis shows that genes carrying L2 in their 3’UTR are present and enriched in AGO2 RIP fractions of human brain tissue samples, suggesting that they are functional miRNA target genes.

When we looked for potential miRNA target sites within the L2 elements in the 3’UTRs of AGO2-bound genes, we found several potential non-canonical target sites with high complementary to the L2-miRNAs: miR-28, miR-95, miR-151a or miR-708 (Figure 4E). To validate the functionality of these non-canonical target sites, we performed luciferase assays and confirmed that all tested target sites are regulated by L2-miRNAs (Figure 4F). Taken together, these data provide functional evidence demonstrating that L2-miRNAs regulate target genes carrying L2-derived sequences in their 3’UTR.

### Dysregulation of L2-microRNAs in human glioblastoma

Since miRNAs and TEs often display aberrant expression in human cancers (Anwar et al. 2017; Banelli et al. 2017), we next investigated whether the expression of L2-miRNAs and their target network are altered in glioblastoma. We first conducted AGO2 RIP followed by small RNA sequencing on human brain biopsies obtained from glioblastoma patients (n = 6). We found that, as in cortex samples, AGO2-bound small RNAs in glioblastoma tissue displayed a high enrichment for 20-24 nt long reads while input samples displayed a broad size profile of RNAs (S2A Fig). When investigating the genomic origin of AGO2-bound small RNAs in glioblastoma, we found very similar results as in cortex tissue with most of the small RNAs being classical miRNAs and only very limited evidence for AGO2-binding of tRFs or piRNA (Fig 5A).

**Figure 5:**
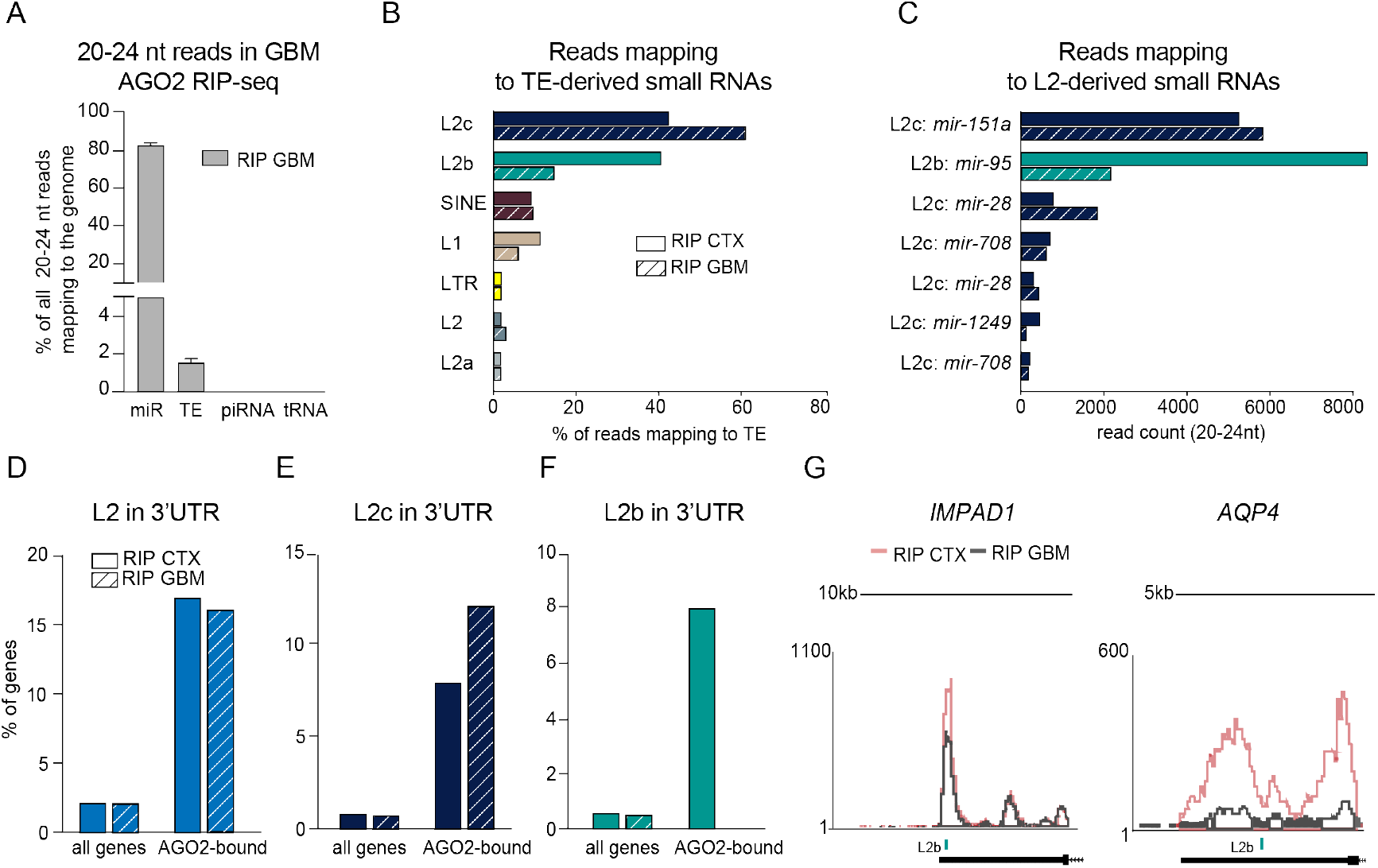
AGO2-associated L2-miRNAs and L2-carrying genes are altered in glioblastoma. A) Bar graph showing the percentage of 20-24 nt long read mapping to mature miRNAs (miR), transposable elements (TE), piwi RNAs (piRNAs) and transfer RNAs (tRNA). Data is represented as mean ± SEM (RIP GBM n = 6). B) Bar graph showing the percentage of 20-24 nt long uniquely aligned reads mapping to transposons in AGO2 RIP samples of cortex (CTX) and glioblastoma (GBM) tissue (RIP CTX n = 3; RIP GBM n = 6). C) Number of uniquely aligned reads mapping to L2 elements giving rise to precursor miRNAs (mir) in AGO2 RIP-seq samples from cortex and glioblastoma tissue (RIP CTX n = 3; RIP GBM n = 6). D-F) Bar plots showing the percentage of genes with L2 / L2b / L2c in the 3’UTR of all genes, and of the top 100 highest expressed genes that are more than 4-fold enriched in RIP compared to input samples of cortex and glioblastoma tissue. G) Overlap of UCSC genome browser tracks showing examples of L2-carrying transcripts with altered AGO2-binding in glioblastoma compared to cortex. AGO – Argonaute, AGO RIP – AGO-RNA Immunoprecipitation, L1 – LINE1, L2 – LINE-2, LTR – long terminal repeat, SINE – short interspersed nuclear element.

We next analysed the miRNA profile in glioblastoma and found miR-21 to be the most abundant miRNA in this tissue (S2B Fig). Differential expression analysis of miRNAs in glioblastoma samples compared to cortex samples showed altered expression of several miRNAs previously identified to be dysregulated in glioblastoma e.g. miR-21, miR-10b and miR-128 (S2C Fig) (Banelli et al. 2017; Masoudi et al. 2018). We then analysed TE-derived miRNAs and found that L2-miRNAs were abundant also in glioblastoma-samples (Fig 5B). Interestingly, when analysing individual L2-miRNAs, we found a much higher proportion of reads mapping to L2b-derived miR-95-3p in cortex than in glioblastoma samples (Fig 5C).

### AGO2 binding of L2-carrying genes is altered in human glioblastoma

To investigate how AGO2-bound genes carrying L2 in their 3’UTR are affected in glioblastoma we conducted AGO2 RIP, followed by total RNA sequencing. We found very similar results to cortex tissue when analysing the global genomic distribution of reads. Input samples showed a high amount of rRNA, which was strongly decreased in the RIP fraction (S2D Fig). Moreover, reads mapping to 3’UTRs of genes and to TEs were enriched in the RIP samples of glioblastoma tissue compared to input fractions (S2D Fig).

We next focused our analysis on transcripts carrying L2 in their 3’UTR and found that while the overall numbers of the respective transcripts were very similar in glioblastoma and cortex samples, there were profound differences when separately analysing L2b and L2c-carrying transcripts (Fig 5E&F). Genes carrying L2c in their 3’UTR were enriched in the RIP samples from both sample types (Fig. 5E). In contrast, L2b-carrying transcripts were more abundant in cortex samples than in glioblastoma tissue, where we did not detect any AGO-bound L2b-carrying genes (Fig 5F&G). This is in line with the small RNA data, where the L2b-derived miR-95 was highly expressed in cortex samples, but not in glioblastoma (Fig 5C). Thus, this analysis directly links the altered expression of L2b-derived miR-95 to the lack of L2b-carrying targets bound to AGO2 in glioblastoma tissue. Interestingly, several of these L2b-targets, which are not bound by AGO2 in glioblastoma, are implicated in glioma, such as *PCDH9* (Wang et al. 2014) and *AQP4* (Lan et al. 2017).

## Discussion

In this study, we demonstrate that L2 elements serve as a source for several miRNAs that are bound by AGO2 proteins in the human brain with the ability to target hundreds of transcripts carrying L2-fragments in their 3’UTR. Our study therefore provides a model for how TEs control post-transcriptional networks in the human brain. L2-miRNAs and targets are ubiquitously expressed across multiple human tissues, generally at low levels. This suggests that this network acts as a post-transcriptional buffer, providing additional robustness to the levels of housekeeping genes. This is in line with previous observations where L2-derived miRNAs, such as miR-95 and miR-151, were found to control basic cellular mechanisms such as lysosomal function and cell migration (Ding et al. 2010; Frankel et al. 2014).

L2 elements invaded the genome of our ancestors some 100-300 million years ago, well before mammalian radiation (Kapitonov 2006). Throughout evolution, these TE sequences likely underwent a high evolutionary pressure, leading to the degradation of non-functional sequences, hence, only leaving fragments of sequences remaining in the genome. Today, a portion of these sequences form a post-transcriptional network composed of miRNAs and miRNA-target templates present in the 3’UTR of thousands of protein-coding genes. It is tempting to speculate that the majority, if not all, miRNAs have originally emerged from TEs, since TEs are scattered throughout the genome in high numbers and therefore have the potential to give rise to a large number of miRNAs and miRNA target sites. However, since most miRNAs are evolutionary old (more than a billion years old) it is impossible to determine their origin with certainty. Several TEs have previously been implicated in both the generation of miRNAs as well as miRNA-targets (Smalheiser and Torvik 2005; Piriyapongsa et al. 2007; Roberts et al. 2013; Boudreau et al. 2014; Roberts et al. 2014). However, most observations concerning TE-derived miRNA-target interaction are based on computational predictions and experimental evidence for TE-derived miRNA-target interaction was up until now limited. Key to the findings in this study is the use of AGO2 RIP on human tissue coupled to next-generation sequencing on both small and total RNA. By using this approach, we could confirm and demonstrate the functionality of L2-miRNAs and show that transcripts with L2 in their 3’UTR are AGO2-bound miRNA targets.

It is commonly accepted that miRNAs regulate genes via the seed sequence, which spans from nucleotides 2-7 in the 5’ end of the miRNA (Bartel 2009). Additionally, several alternatives to canonical seed sequences have been proposed (Shin et al. 2010; Kim et al. 2016). For instance, miR-151 and miR-28 have previously been found to use centred seed pairing to regulate genes (Shin et al. 2010). Since TE-derived miRNAs show high sequence similarities with TE sequences in the 3’UTR of genes, such miRNAs can most likely use extensive base-paring for target recognition and regulation. This suggests that non-canonical miRNA target sites might be more broadly used than previously thought.

Glioblastoma is the most aggressive and most common type of brain tumour in adults and is characterised by vast cellular and genetic heterogeneity. In the material used in this study, we found that the previously identified glioma-associated miRNAs such as miR-21 and miR-10b were dysregulated. In addition, we found that the L2b-derived miR-95 was highly expressed in cortex, but not in GBM, which is in line with previous reports demonstrating reduced levels of miR-95 in glioma (Skalsky and Cullen 2011; Fan et al. 2015). Together with our finding that L2b-carrying transcripts are not bound to AGO2 in glioblastoma tissue compared to cortex samples, this suggests that miR-95 could play an important role in the regulation of transcripts related to tumour progression or tumour defence in glioblastoma. Future studies analysing the function of miR-95 in glioblastoma are therefore warranted.

In summary, this work demonstrates the existence of a TE-based post-transcriptional regulatory network that shapes the expression of hundreds of TE-carrying transcripts and hence provides an additional mechanism for TEs to influence crucial gene networks human tissues.

## Methods

### Human Tissue

Fresh human adult neocortical and glioblastoma tissues were obtained during resective surgery in patients suffering from glioblastoma or pharmacologically intractable epilepsy, respectively. The tissue was snap frozen immediately following removal. The use of human brain tissue was approved by the local Ethical Committee in Lund (212/2007 for epilepsy and H15 642/2008 for GBM) in accordance with the declaration of Helsinki. Prior to each surgery written informed consent were obtained from all subjects.

### Mouse brain tissue

All animal-related procedures were approved and conducted in accordance with the committee for use of laboratory animals at Lund University. For AGO2 RIP-seq, the striata of mice were quickly dissected and immediately homogenised and lysed in ice-cold lysis buffer.

### AGO2 RIP-seq

Fresh or snap-frozen tissue was homogenised in ice-cold lysis buffer (10 mM HEPES (pH = 7.3), 100 mM KCl, 0.5 % NP40, 5 mM MgCl_2_, 0.5 mM dithiothreitol, protease inhibitors, recombinant RNase inhibitors, 1 mM PMSF) using TissueLyser LT (30 Hz, 4 minutes).

Homogenates were centrifuged twice for 15 minutes at 16,200 x g, 4 °C to clear the lysate. 1/10 of the sample was then saved as input (total RNA) control. The remaining lysate was incubated with anti-AGO2-coated Dynabeads^®^ Protein G beads (Life Technologies) at 4 °C for 24 hours with end-over-end rotation (AGO2 antibody: anti-Ago2-3148 for human brain tissue (Grey et al. 2010) and Sigma-Aldrich 2E12-1C9 for mouse tissue).

After incubation, the beads were collected on a Dynamagnet (1 minute, on ice) and gently resuspended in low-salt NT2 buffer (50 mM Tris-HCL (pH = 7.5), 1 mM MgCl_2_, 150 mM NaCl, 0.5 % NP40, 0.5 mM dithiothreitol, 1 mM PMSF, protease inhibitors and recombinant RNAse inhibitors). The beads were transferred into a new collection tube and washed once with low-salt NT2 buffer, followed by two washes with high-salt NT2 buffer (50 mM Tris-HCl (pH = 7.5), 1 mM MgCl2, 600 mM NaCl, 0.5 % NP40, 0.5 mM DTT, protease inhibitors, 1 mM PMSF and recombinant RNAse inhibitors). After the last washing step, the RNA fraction was resuspended in QIAzol buffer and RNA was isolated from RIP and input samples according to the miRNeasy micro kit (Qiagen).

### cDNA library preparation and sequencing of human AGO2 RIP samples

cDNA library preparation was conducted using the NEB small RNA library prep kit for small RNA sequencing and the NuGen Ovation RNAseq V2 kit, followed by the Ovation^®^ Ultralow V2 Library or Ovation^®^ Rapid Library Systems, for total RNA sequencing. Illumina high-throughput sequencing (HiSeq2500 SR 1×50 run and HiSeq3000 1×50) was applied to the samples (total number of reads for small RNA sequencing: 344508344; total number of reads for total RNA sequencing: 527413532) at the UCLA Microarray Core Facility.

### Analysis of the genomic distribution of small RNA sequencing reads from AGO2 RIP-seq

For the analyses of the genomic distribution of reads, the data was aligned to the human reference genome (hg38) using STAR (2.5.0a) (Dobin et al. 2013) allowing multimapping and two mismatches per 22bp (--outFilterMismatchNoverLmax 0.05).

Reads were quantified with the SubRead package FeatureCounts (Liao et al. 2014) (minimal overlap 19 nt) using annotations obtained from miRbase (Kozomara and Griffiths-Jones 2014), UCSC genome browser RepeatMasker track (GRCh38) and piRNAbank (Sai Lakshmi and Agrawal 2008).

### Analyses of TE-derived small RNAs bound to AGO2

For the analyses of small RNAs derived from individual TEs, the data was aligned to the human reference genome (hg38) using STAR (2.5.0a), allowing for 0 mismatches. Reads that mapped equally well to more than one locus were discarded (-- outFilterMultimapMax 1), while reads with a better alignment score for one locus than for any other position were kept (--outFilterMultimapScoreRange 1). These parameters allow for the quantification of the specific, expressed locus, while reads with repeated identical sequences that cannot be assigned to any specific locus were discarded. As control, multimapping reads were kept, however there were no significant effects on the quantification and detection of TE-derived small RNAs, indicating that there are no or very few identical TE-derived smallRNAs.

For detecting TE-derived miRNAs, only reads with a length of 20-24 nt were included in the analysis. Furthermore, for the identification of high-confidence miRNAs, 3p and 5p strands and typical Dicer cleavage patterns had to be present. We also assessed conservation of the small RNAs by using the 100 vertebrates Basewise Conservation data by phyloP from the PHAST package.

For the alignment of the L2c-derived small RNAs to the L2c consensus, we used the raw alignment of the L2c consensus to the reference genome (GRCh38) generated by RepeatMasker to annotate genomic L2c. This alignment can be assessed through the detailed visualisation of RepeatMasker annotations in the UCSC genome browser. The genomic-to-L2c consensus alignment by RepeatMasker was used to map the TE-derived small RNA to its relative location in the L2c consensus.

### Expression analysis of L2-small RNAs in different tissues

To analyse the expression of L2-small RNAs, we downloaded expression profile data of miRNAs in 400 human samples from http://fantom.gsc.riken.jp/5/suppl/De_Rie_et_al_2017/ (file: human.srna.cpm.txt) (de Rie et al. 2017) and extracted the counts per million (cpm) data for the 19 L2-miRNAs, we identified in this study (Table 1). A pseudo-count of 1.0 was added and the logarithm (base 2) was taken for the values. Hierarchical clustering was done using Euclidean distance with single linkage.

### Expression analysis of genes carrying L2 in 3’UTR

For obtaining a list of transcripts with L2 elements in their 3’UTRs, Ensembl 3’UTR annotations were overlapped with L2 RepeatMasker annotations using BEDTools intersection requiring at least 1 bp overlap. A publicly available dataset consisting of transcriptome levels of 27 different tissue types across 95 individuals was downloaded from www.ebi.ac.uk/arrayexpress/ (accession ID E-MTAB-1733) (Fagerberg et al. 2014). FPKM values for 1932 genes out of the 2042 genes previously identified as having L2 elements in their 3’UTRs were extracted from the above data. These genes were grouped into seven different classes depending on their tissue-specificity characteristics as described in Fagerberg et al. (Fagerberg et al. 2014). A pseudo-count of 1.0 was added and logarithm (base 2) was computed before performing hierarchical clustering (Euclidean distance and single linkage).

### Analysis of structural conservation of L2 elements in human genome

To get the coverage of bases derived from each base position in L2c consensus, the relative position of genomic L2c elements aligning to L2c consensus were extracted from the RepeatMasker output file (http://hgdownloadtest.cse.ucsc.edu/goldenPath/hg38/bigZips/hg38.fa.out.gz). The coverage module of BEDTools (v2.24.0) was used to calculate the consensus coverage, which shows how many of the genomic L2c copies have the given position in the consensus sequence.

### Analyses of total RNA sequencing data from AGO RIP on human tissue

For the expression analyses of AGO2-bound genes, reads were aligned to the human reference genome (hg38) using STAR (2.5.0a) with default settings. 3’UTR annotations were obtained from Ensembl (Kersey et al. 2016).

### Analyses of total RNA sequencing data to identify L2-carrying transcripts

For the identification of L2 elements within 3’UTRs, reads were aligned to the human reference genome (hg38) using STAR (2.5.0a), allowing for 2 mismatches per 50 nt (--outFilterMismatchNoverLmax 0.04), and discarding multi-mapping reads. The obtained files were intersected with L2 annotations, obtained from RepeatMasker with the requirement of 80 % overlap (-f 0.8). The reads were then intersected with 3’UTR annotations (Ensembl) with a requirement of at least 1 bp overlap. The intersection was conducted using the BEDTools (2.26.0) intersect module.

### Identification of miRNA target sites in L2 elements in the 3’UTR of genes

The L2 sequence in the 3’UTR of a gene was aligned to the sequence of a L2-miRNA using the EMBOSS Needle nucleotide pairwise alignment tool (Rice et al. 2000) to find sequence complementarity. Following parameters were used: DNAfull substitution scoring matrix, 10 gap open penalty and 0.5 gap extend penalty, no end gap penalty applied.

### Luciferase reporter assay

A 400 bp sequence incorporating the L2-derived miRNA binding sites in the 3’UTR of *CLIP3*, *IMPAD1*, *YWHAZ*, *TPP1* and *ADD2* was cloned into the dual luciferase reporter vector pSICHECK-2 (Promega). The luciferase reporter constructs were co-transfected with either a GFP overexpression vector or the respective L2-derived miRNA overexpression vector into three independent replicates of 293T cells using Turbofect (Fermentas). 48 hours after transfection, cells were assayed for luminescence using a dual-luciferase assay (Promega). One-way ANOVA followed by a Tukey’s multiple comparison post hoc test were performed in order to test for statistical significance. Data is presented as mean ± SEM.

## Data access

All RNA sequencing data have been submitted to NCBI Gene Expression Omnibus database and assigned the GEO series accession number GSE106810.

## Acknowledgement

We would like to thank Stefan Thor, Volker Busskamp and Didier Trono as well as all members of the Jakobsson lab for useful comments on the manuscript. We also thank J. Johansson, M. Persson Vejgården, U. Jarl, A. Hammarberg, E. Ling, B. Mattson, S. da Rocha Baez and M. Sparrenius for technical assistance.

The work was supported by grants from the Swedish Research Council, the Swedish Foundation for Strategic Research, the Swedish Brain Foundation; the Swedish excellence project Basal Ganglia Disorders Linnaeus Consortium (Bagadilico), and the Swedish Government Initiative for Strategic Research Areas (MultiPark & StemTherapy).

## Author contributions

R.P., P.L.B, Y.S., M.J., K.P., J.B. and J.J. designed research and analyzed data. R.P., M.J., K.P. performed research. R.P., P.L.B. and Y.S. performed Bioinformatics analysis. R.P. and J.J. wrote the paper and all authors reviewed the manuscript.

---

**Supplemental Fig 1: Related to Figure 1.**
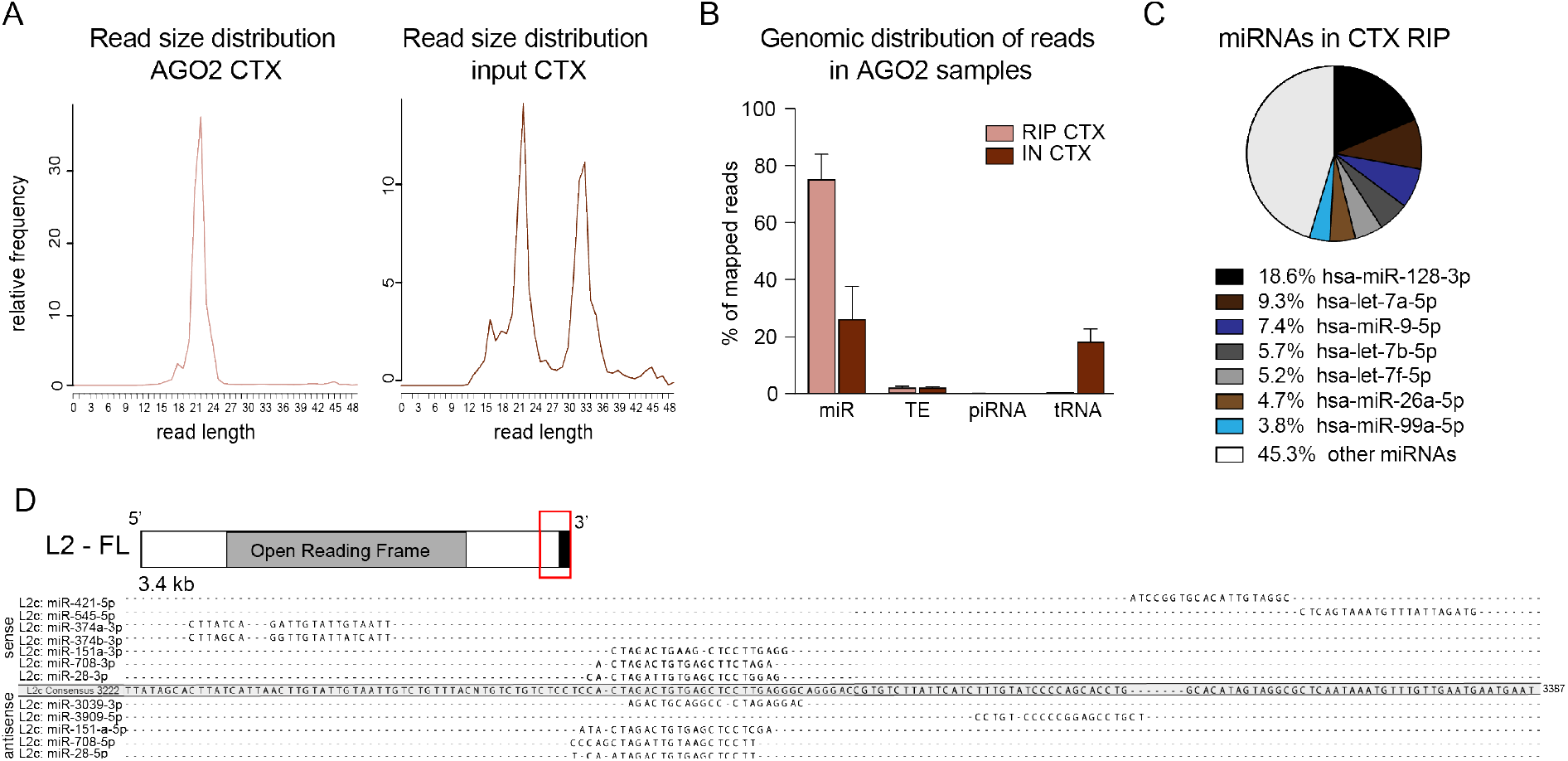
A) Read size distribution of cortex (CTX) RIP and input samples. B) Bar graph showing the percentage of 20-24 nt long reads in cortex tissue mapping to mature miRNAs (miR), transposable elements (TE), piwi RNAs (piRNA) and transfer RNAs (tRNA). Data is represented as mean ± SEM (RIP CTX n = 3). C) Pie chart showing the percentage of reads mapping to individual miRNAs. D) Schematics of a full-length (FL) L2 element (red box indicates the 3’ end) and alignment of L2c-miRNAs to the 3’ end (position 3222 – 3387) of the L2 consensus sequence. AGO - Argonaute.

**Supplemental Fig 2: Related to Figure 5.**
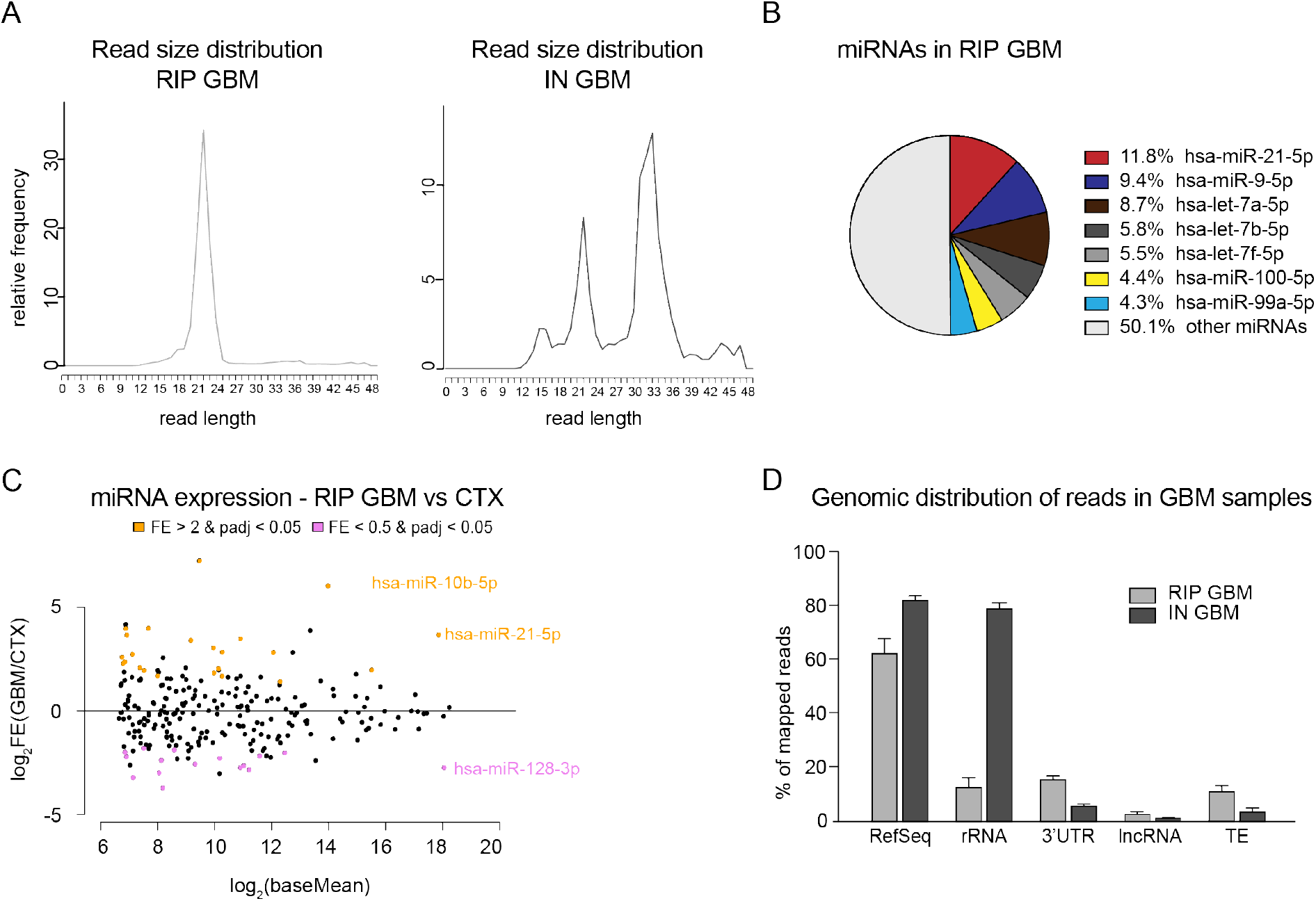
A) Read size distribution of glioblastoma RIP and input samples. B) Pie chart showing the percentage of reads mapping to individual miRNAs in glioblastoma tissue. C) Dot plot depicting miRNA expression in glioblastoma RIP compared to cortex RIP samples. The log2 transformed base Mean is plotted against the log2 transformed fold change. BH-adjusted p value < 0.05. D) Genomic distribution of total RNA sequencing reads in AGO2 RIP and input samples (GBM, n = 5). Data is presented as mean ± SEM. E) Table of genes with L2b in their 3’UTR that show altered AGO2 binding in glioblastoma tissue. AGO – Argonaute, FE – fold enrichement, GBM – glioblastoma.

